# Functional characterization of the lateral septal calbindin neurons in maternal care

**DOI:** 10.64898/2026.02.11.705266

**Authors:** Vivien Szendi, Gina Puska, Máté Egyed, Miklós Márton Takács, Szilvia Bartók, Júlia Puskás, Petra Varró, Attila Szűcs, Árpád Dobolyi

**Author notes:** The first two authors contributed to the work equally. Corresponding author: Dr. Árpád Dobolyi, Department of Physiology and Neurobiology, Eötvös Loránd University, Budapest, Hungary. /8775.

## Abstract

The ventral subdivision of the lateral septum (LSv) is a forebrain region linked to maternal care with a high density of calbindin-containing (Cb^+^) neurons whose properties, connections and functions remain unknown. In the present study, it was established that the majority of pup-activated neurons are inhibitory Cb^+^ in the LSv and the density of activated neurons in the LSv is close to that of the medial preoptic area (MPOA), a central brain area in maternal care regulation. Electrophysiological recordings of LSv^Cb+^ and LSv^Cb-^ neurons revealed that LSv^Cb+^ neurons exhibited higher membrane resistance, lower spike amplitude and rise slope compared to LSv^Cb-^ neurons. The chemogenetic inhibition of LSv^Cb+^ neurons led to reduction in pup-licking behavior without affecting other maternal behavior or eliciting anxiety- or depression-like behavior. To regulate licking behavior, LSv^Cb+^ neurons connect with other maternally involved brain areas. Anterograde and retrograde tract-tracing revealed that two-thirds of the MPOA-projecting LSv neurons are Cb^+^. A subset of neurons within the posterior intralaminar nucleus (PIL) in the lateral thalamus, a proposed relay nucleus of tactile and auditory information from the pups, express a maternally induced neuropeptide, the parathyroid hormone 2 (PTH2). Their fibers closely apposed Cb^+^ neurons in the LSv. Using double labeling, we also identified PTH2 receptors on LSv^Cb+^ neurons, which are presumably activated by PTH2 released from nearby terminals. Using electron microscopy, we confirmed synaptic connection between PTH2+ fibers and inhibitory neurons in the LSv. The results provide evidence that Cb^+^ neurons of the LSv are components of the maternal circuitry.

**Significance Statement:** Maternal care depends on pup cues, but many responsible circuit elements remain uncharacterized. We identify inhibitory calbindin-expressing neurons in the ventral subdivision of the lateral septum (LSv) containing a major pup-activated neuronal population with distinctive electrical properties. Their selective inhibition reduced pup-licking without affecting other maternal, anxiety- or depression-like behaviors, suggesting a specific role in caregiving. Detailed circuit tracing data revealed that a significant proportion of LSv calbindin-positive neurons extend their projections to the medial preoptic area, a well-established core maternal regulatory region. We further demonstrate synaptic input from maternally relevant parathyroid hormone 2 (PTH2)-expressing thalamic afferents onto LSv inhibitory neurons, possibly conveying pup stimuli. The data suggest that LSv calbindin neurons are key nodes in maternal circuitry.

## Introduction

Well-being and survival of mammalian offspring are highly dependent on the appropriate caregiving behavior of their mothers. In the absence of proper maternal care, offspring may develop long-term behavioral and physiological disorders (Avraham et al., 2020; Sacks et al., 2016; Taka-Eilola et al., 2019), therefore, understanding the neural regulation of maternal behavior is crucial for developing effective clinical interventions.

Over the years, several brain regions have been implicated in the regulation of the maternal state, including the lateral septum (LS). LS is a subcortical forebrain structure that integrates contextual and social environmental information to guide motivated behaviors. It participates in the regulation of a wide range of social and non-social behaviors through its anatomically and functionally distinct subregions, diverse neuronal populations, and dense reciprocal connections with multiple brain areas (Leroy et al., 2018; Rashid et al., 2024; Risold & Swanson, 1997b; Yeates et al., 2022). Lesions involving the LS cause disruption in different aspects of maternal care including nursing and pup retrieval (Cruz & Beyer, 1972; Fleischer & Slotnick, 1978; Olazábal et al., 2013). Indeed, pup-induced activation of LS is also prominent in rat and rabbit (Aguirre et al., 2017; González-Mariscal & Poindron, 2002; Lonstein et al., 1997). Furthermore, LS neurons converge hippocampal and amygdalar inputs (Isaac et al., 2025; Risold & Swanson, 1997b), that are important in recognizing and remembering pup olfactory cues ensuring proper maternal care (Abellán-Álvaro et al., 2022), maternally relevant hormones, peptides, and pup-originated cues into maternal behaviors (Puska et al., 2025). These signals are processed by distinct populations of hormone-receptor-expressing neurons, including oxytocin receptor (OTR)–positive cells. Oxytocin is a key modulator of maternal behavior, and OTR-expressing LS neurons have been shown to regulate social fear. Notably, lactating female mice display reduced social fear, an effect that is abolished when OTR-expressing LS neurons are inhibited (Menon et al., 2018).

For successful maternal care, cues originating from the pups must be accurately detected, integrated, and translated into appropriate behavioral outputs, a function for which the LS is well suited as a central hub. Recently, our group identified a maternally relevant circuit involving the LS that receives dense input from neurons expressing parathyroid hormone 2 (PTH2), a neuropeptide strongly induced during lactation, in rat dams (Puska et al., 2025). PTH2 neurons are located in the posterior intralaminar nucleus (PIL) of the lateral thalamus (Yang et al., 2025). Previous studies demonstrated that expression of PTH2, also known as tuberoinfundibular peptide 39 (TIP39), increases dramatically after parturition and peaks around postpartum day 9 (Cservenák et al., 2013). We further showed that the ventral subdivision of the LS (LSv) contains a population of pup-exposure-activated, c-Fos-expressing neurons whose density is comparable to that observed in the medial preoptic area (MPOA) and the paraventricular hypothalamic nucleus (PVN) (Puska et al., 2025). Both area are part of the hardwired maternal circuitry (Kuroda et al., 2024), especially MPOA, where neuronal activity is essential for the initiation, maintenance, and fine regulation of multiple components of maternal behavior. Based on the robust pup-induced c-Fos activation in the LSv, the present study aimed to identify the specific subpopulation of LSv neurons activated in the maternal context, to characterize their electrophysiological properties, and to determine their functional contribution to maternal care. Although previously distinguishable tonic (regular spiking), phasic (single spiking) and burst firing cells were recorded from the LSv (Wang et al., 2019), cell-specific electrophysiological characterization is not defined in the area.

Moreover, we previously found that LSv neurons activated in rat dams project prominently to the MPOA. Therefore, in the current study, we also sought to examine whether this maternally activated LSv subpopulation forms a functional component of the neural circuitry that directly executes and modulates maternal behavior.

## Material and methods

### Animals

All procedures involving mice were executed according to the experimental protocols that meet the guidelines of the Animal Hygiene and Food Control Department, Ministry of Agriculture, Hungary (40/2013) in accordance with EU Directive 2010/63/EU for animal experiments. This study was approved by The Workplace Animal Welfare Committee of the National Scientific Ethical Committee on Animal Experimentation at Eötvös Loránd University, Budapest (PE/EA/117-8/2018, PE/EA/568-5/2021, PE/EA/00086-6/2025).

A total of 43 experimental animals were used in the study. 12 C57BL/6Ncrl (Charles Rivers Laboratories, Hungary, https://www.criver.com/products-services/find-model/c57bl6-mouse?region=3631) female mother mice were used to determine maternal c-Fos-activation patterns. During tissue preparation, 3 glasses of parallel brain slices were cut, thus, three different immunohistochemical procedures could be done on sections that originated from the same animals. In this case the slices from the 12 mice were used for peroxidase-based c-Fos immunohistochemistry, for calbindin/c-Fos double fluorescent immunolabeling, and for histology to examine pup dependent activation of PTH2-expressing neurons in the PIL (in all cases: pup-induced group: n = 6, pup-deprived control group: n = 6). Additional 6 C57BL/6Ncrl female mother mice and 2 Wistar (Charles Rivers Laboratories, Hungary, https://www.criver.com/products-services/find-model/wistar-igs-rat?region=3631) mother rats were used to immunolabel and analyze PTH2+ fibers around maternally activated Cb^+^ neurons in the LSv.

A total of 12 Calb1-2A-dgCre-D (Cb-Cre, The Jackson Laboratory, https://www.jax.org/strain/023531, RRID: IMSR_JAX:023531) virgin female mice were used for this study. 10 of them were involved in chemogenetic manipulation of LSv^Cb+^ neurons (inhibition: n = 5, control: n = 5), and 2 were used for cell type specific anterograde tracing, while an additional 2 C57BL/6NCrl female mice were required for the retrograde tract-tracing experiment. All of the animals involved in stereotaxic surgery were 8 weeks old on the day of the surgery and they were allowed to recover for three weeks after the procedure before further testing or sacrificing. After the surgery, animals were individually housed.

For electrophysiological measurements on brain slices, 5 Cb-ZsGreen female mice were sacrificed, which are the offspring of Calb1-2A-dgCre-D and GtROSA26Sor_CAG/ZsGreen1 (The Jackson Laboratory, https://www.jax.org/strain/007906, RRID: IMSR_JAX:007906) mice. In these animals, Cb^+^ neurons express the ZsGreen fluorescent protein, so that the recognition of these neurons is possible in the patch clamp setup equipped with fluorescent microscope. 3 additional Cb-ZsGreen female mice were used for calbindin histochemistry.

For correlative light and electron microscopy (CLEM), one VGAT-ZsGreen mother mouse was used, which is a crossing of VGAT-IRES-Cre (The Jackson Laboratory, https://www.jax.org/strain/016962, RRID: IMSR_JAX:016962) and GtROSA26Sor_CAG/ZsGreen1 mice.

When mother animals were used, females were mated with male conspecifics from the same strain, then kept individually housed and the number of pups was balanced to 8 within 3 days of delivery and they were sacrificed on the 9th postpartum day to ensure proper PTH2 expression for histological experiments.

If not stated otherwise, animals were kept group housed under standard laboratory conditions of 12/12 light-dark cycle with light on at 6.00 am and food and water were accessible ad libitum. Standard polycarbonate cages were used for keeping the animals and for home-cage testing.

### Pup-deprivation protocol used for c-Fos studies

To determine the c-Fos-activation pattern in their brains, mother mice were deprived from their pups for 20 hours on postpartum day 8-9. Separation was inevitable to establish basal c-Fos-activation level hence control mothers were not reunited with their pups and were sacrificed after deprivation time to determine the standard activation pattern during analysis. Following deprivation, mothers received their pups back for 2 hours during which they could freely interact with them including suckling and were sacrificed after that as c-Fos expression shows the highest level at 1.5-2 hours from reintroducing the pups. A total of 12 mice were used in this experiment.

### Electrophysiological recordings

To characterize the electrophysiological properties of lateral septum neurons, we utilized whole-cell patch clamp and applied current step protocols to elicit their sub- and suprathreshold voltage responses. Here, coronal acute brain slices of 250 µm thickness were prepared using a vibratome (Electron Microscopy Sciences) and incubated in room temperature artificial cerebrospinal fluid (ACSF) for a minimum of 1 hour before patch clamp recordings were initiated. The composition of ACSF was (in mM): NaCl (126), NaHCO_3_ (26), KCl (1.8), KH_2_PO_4_ (1.25), CaCl_2_·2H_2_O (2.4), MgSO_4_·7H_2_O (1.3), D-glucose (10). The patch electrodes were pulled from borosilicate glass, had 6-8 MOhm resistance and were filled with the following intracellular solution (in mM): K-gluconate (100), KCl (10), KOH (10), MgCl2 (2), NaCl (2), HEPES (10), EGTA (0.2), D-glucose (5); the pH was set to 7.35. To stimulate neurons and record their voltage responses we used a Multiclamp 700B (Molecular Devices) amplifier and the DASYLab 11 (National Instruments) data acquisition program. Our main stimulus protocol was designed to gain an accurate estimation of the cells’ many physiological parameters and measure their intrinsic excitability. Here, we stimulated the neurons using current step injections of 400 ms duration in 1.25 s cycles, starting at −80 pA and incremented by 4 pA until robust firing of the neuron was observed. For each negative current step, membrane resistance, time constant, and voltage sag values were calculated and plotted as functions of the injected current. Linear fitting of such relationships and extrapolation was used to calculate resting levels of membrane resistance, voltage sag index, and membrane time constant at I=0 pA. All the calculations were performed by the NeuroExpress analysis suite developed by ASz.

### Stereotaxic surgery for chemogenetic manipulation and neuronal tracing

For chemogenetic silencing of the LSv^Cb+^ neurons, anterogradely spreading rAAV-hSyn-DIO-hM4D(Gi)-mCherry (a gift from Bryan Roth, Addgene plasmid # 50475; http://n2t.net/addgene:50475; RRID: Addgene_50475) and for controls rAAV-hSyn-DIO-mCherry (a gift from Bryan Roth, Addgene plasmid # 50459; http://n2t.net/addgene:50459; RRID: Addgene_50459) viral vectors were injected into the LSv of virgin female Cb-Cre mice (inhibition: n = 5, control: n = 5). As a result of the used transgenic mouse strain and the construct of the viral vector, mCherry and the modified inhibitory human muscarinic receptor (hM4D(Gi)) were only expressed in the Cb^+^ neurons. To define the projections of LSv^Cb+^ neurons, the control virus used in chemogenetic experiments (rAAV-hSyn-DIO-mCherry) was injected unilaterally into the LSv of Cb-Cre mice (n = 2). For tracing and examining the input from LSv arriving to the MPOA, retrogradely transporting cholera toxin beta subunit (CTB, List Biological Laboratories, Campbell, CA) was injected into the MPOA of female C57BL/6NCrl mice (n = 2).

First, animals were anesthetized using ketamine-xylazine solution (90-100 mg/body weight kg) then were placed into the stereotaxic apparatus. Following the shaving and disinfection of the head, a hole of 1.5 mm diameter was drilled into the skull above the injection coordinates. The used stereotaxic coordinates (Paxinos & Franklin, 2021) were the following for LSv: AP: +0.5; LM: +/- 0.5; DV: −3.15 and for the MPOA: AP: +0.05; LM: −0.25; DV: - 5.2. For injection, glass capillaries (World Precision Instrument) with 1.14 mm diameter were filled first with mineral oil (Sigma-Aldrich) then placed into the Nanoinjector apparatus (Nanoinjector 2010, World Precision Instrument) followed by 1 µm of virus. For chemogenetic manipulation bilaterally while for neuronal tracing unilaterally, 30 nl of the above-mentioned viral vector and 50 nl of CTB were targeted into the LSv and MPOA, respectively at a speed of 50 nl/min. The capillary was left in place for 5 minutes then slowly removed. Animals were left to recover for three weeks before the behavioral tests or sacrifice began.

### Clozapine-N-oxide (CNO) injection

For chemogenetic manipulation, 45 minutes before the test started, clozapine-N-oxide (CNO) or vehicle solution was injected intraperitoneally. CNO is the exogenous ligand of the used modified receptor and is dissolved in 5% dimethyl sulfoxide (DMSO) in distilled water solution. On the first and second control days, the animals received only DMSO solution (1 ml/kg). On test day, CNO solution in a 0.3 mg/kg dose was injected into the mice.

### Measurement of spontaneous maternal behavior

Viral injected virgin females were sensitized by receiving pups for 10 minutes for two consecutive days before the test started. During the sensitization period, mice could freely interact with the pups, but the interaction was supervised by the research leader to avoid attacks toward the pups. No aggressive behavior toward pups was seen.

Experimental protocol for spontaneous maternal behavior test was designed to be self-controlled meaning that the animals were examined on both control and test days, and the behaviors of the same animal on different days were compared. There were two control days, one preceding the test day and one two days after the test day on which animals received an intraperitoneal injection of the vehicle solution, while on the test day they received CNO solution in similar manner.

Throughout the spontaneous maternal behavior test, three pups were placed in three corners of the home cage of the test animal while the test animal was placed in the fourth corner where the nest was located. During the test, examined mice could freely interact with the pups under video recording for 30 minutes, from which 20 minutes were evaluated manually using the Solomon Coder open-source software (https://solomon.andraspeter.com/). Four different behavior elements were examined namely ‘*pup sniffing*’ meaning that the examined mouse sniffed the body or head of a pup, ‘*pup grooming*’ that is the extensive grooming and nursing of a pup by the examined mouse, ‘*pup licking*’ described as the licking of the body or the anogenital region of pups by the examined mouse, and ‘*nest building*’ during which the mouse built or refined the nest material. The time spent with each element (duration) was examined.

### Testing of anxiety- and depression like behavior

On the CNO injection day, following spontaneous maternal behavior test, additional tests were executed to determine if possible maternal behavioral changes are caused by other factors. To examine this, open field, elevated plus maze (EPM), and forced swim tests were used.

During the open field test, test animals were placed into the open field apparatus (40 x 40 cm) for 20 minutes under video surveillance. Recordings were evaluated automatically using SMART Video Tracking software v3.0 (Panlab Harvard Apparatus) by analyzing the time spent in the corners of the apparatus, next to the walls of the apparatus, and in the center of the apparatus, and the total distance travelled by the animal. The EPM apparatus has four arms located 0.5 m above ground from which two have walls (closed arms) while the other two lack them (open arms). Animals’ behavior in the EPM apparatus was recorded by a camera for 10 minutes and videos were analyzed manually using Solomon Coder. Time spent in the closed and open arms and in the area among the four arms (center) was examined. Both of these tests assess the anxiety level of the animals. We examined whether chemogenetically manipulated animals spent more time around corners, walls, and in the closed arms compared to control mice meaning an increased level of anxiety which could influence maternal behavior, as well. Forced swim test (FST) was used to ascertain depression-like state of the animal by placing the examined mouse into a transparent inescapable glass tank filled with 15 cm of 23-24 °C water for 6 minutes. Time spent climbing (escaping-like behavior during which the mice try to climb out of the tank), swimming (the animal is actively swimming in the tank), and floating (depression-like state, only movements necessary to remain above water can be seen) are measured using Solomon Coder. The behavior of chemogenetically manipulated mice was compared to control mice to determine depression-like alteration in behavior.

### Sample preparation for immunohistochemistry

Ketamine/xylazine-hydrochloride (90-100 mg/body weight kg) diluted in 0.9% saline was used for deep anesthesia before perfusion. The transcardial perfusion with 0.9% saline followed by 4% paraformaldehyde solution dissolved in phosphate buffer (PB, pH 7.2) was applied. Then, 50 μm-thick sections were prepared from the brains using a vibratome (VT1000S, Leica).

### Peroxidase-based c-Fos, mCherry and PTH2R immunohistochemistry

To block endogenous hydrogen-peroxidase activity, 0.1% hydrogen-peroxide (H_2_O_2_) containing PB was applied to every third free-floating section for 15 min, then incubation in blocking PB consisting of 1% bovine serum albumin (BSA), 10% fetal bovine serum (FBS), and 0.5% TritonX-100 happened for an hour to block non-specific binding sites. This was followed by the overnight application of c-Fos primary antibody (1:5000, rabbit, Abcam, Cat. #: ab190289), mCherry primary antibody (1:3000, chicken, EnCore Biotechnology Inc., Cat. #: CPCA-mCherry) or anti-PTH2R primary antibody was applied (1:5000, rabbit, a gift from Ted B. Usdin) diluted in PB containing 0.5% BSA, 5% FBS and 0.25% TritonX-100. After washing steps, sections were treated with biotinylated secondary antibody (1:500, donkey anti-rabbit, Jackson Immunoresearch) for an hour followed by an hour-long incubation with avidin-biotin complex (ABC) solution (1:1000, Vector Laboratories). Signal was visualized using 0.1% 3,3’-diamino-benzidine (DAB) solution in Tris buffer (pH 7.4) enhanced by nickel-(II)-sulfate triggered by 0.01% H_2_O_2_ for 7 minutes. Slices were mounted by DePeX mounting medium (Sigma-Aldrich).

### Fluorescent immunohistochemistry

For fluorescent labelling, sections were incubated in the aforementioned blocking PB solution for an hour followed by the application of the proper primary antibody solution in 0.5% BSA, 5% FBS and 0.25% TritonX-100 PB. For calbindin/c-Fos and calbindin/PTH2R double labeling anti-calbindin (1:2000, mouse, Synaptic Systems, Cat. #: 214 011) and anti-c-Fos (1:5000, rabbit, Abcam, Cat. #: ab190289) or anti-PTH2R (1:5000, rabbit, a gift from Ted B. Usdin) were used one-by-one on two consecutive days then fluorescent secondary antibodies of Alexa Fluor 488 (A488, 1:500, donkey anti-mouse for calbindin/c-Fos and anti-rabbit for calbindin/PTH2R, Jackson Immunoresearch) and Alexa 594 (A594, 1:500, donkey anti-rabbit for calbindin/c-Fos and anti-mouse for calbindin/PTH2R, Jackson) were applied for 1.5 hours, respectively. For calbindin/mCherry immunohistochemistry, the anti-calbindin antibody (1:3000, rabbit, Swant, Cat. #: CB38) was applied followed by the incubation of anti-mCherry antibody (1:3000, chicken, EnCore Biotechnology Inc., Cat. #: CPCA-mCherry) on the next day. As secondary antibodies, A488 (1:500, donkey anti-rabbit, Jackson) and A594 (1:500, donkey anti-chicken, Jackson) were applied for 1.5 hours, finished with 10 minutes of incubation in DAPI solution (1:8000, Merck). The same mCherry antibody, anti-chicken secondary antibody, and DAPI were used for fluorescent mCherry labeling for anterograde tracing. To label calbindin on brain slices from VGAT-ZsGreen mice, the aforementioned rabbit anti-calbindin antibody was used. For double labeling calbindin and cholera toxin B subunit (CTB), sections were incubated in the rabbit anti-calbindin antibody solution, then in anti-CTB primer antibody solution (1:10000, goat, List Biological Laboratories, #703). Signals were visualized using anti-mouse A594 and A488 (1:500, anti-goat, Jackson), while for nuclei detection DAPI solution was used as mentioned above. For PTH2/c-Fos histology, anti-PTH2 primary antibody (1:3000, rabbit, Research resource identifier, AB_1235466) was applied for 48 hours followed by the treatment with anti-c-Fos primary antibody (1:15000, guinea pig, Synaptic Systems, Cat. #: 226 004). In this case anti-rabbit A488 and A594 (1:500, donkey anti-guinea pig, Jackson) were used as secondary antibodies. Finally, for triple immunolabeling of PTH2, calbindin and c-Fos, first, the rabbit anti-PTH2 antibody was applied for 48 hours, then mouse anti-calbindin and guinea pig c-Fos antibody were used overnight. Signals were visualized using anti-rabbit A488, DyLight 549 (1:500, donkey anti-mouse, Vector Laboratories) and Alexa Fluor 647 (1:500, donkey anti-guinea pig, Jackson), respectively.

During electrophysiological recording of calbindin neurons, some cells were filled up with biocytin, which were later visualized to determine morphological traits. For visualization, slices were treated with 1% TritonX-100 PB solution overnight, then with 20% BSA and 0.3% TritonX-100 PB for an hour. Finally, streptavidin-TRITC (1:300) solution was applied for 3 hours then washed before mounting the sample.

For every fluorescent immunolabeling, sections were mounted by Aqua-Poly/Mount mounting medium (Polysciences Inc.)

### Light microscopy and image acquisition

Peroxidase-based c-Fos immunolabeled samples were examined using a Nikon Eclipse Ci-L microscope mounted with a Tucsen Michrom 20 Camera. Images were acquired using 4 x (CFI Plan Fluor 4X, N.A. 0.13, W.D. 17.1 mm) and 20 x (CFI Plan Achromat DL 20X N.A. 0.40) objectives. Image adjustments, including brightness/contrast and exposure were created by Adobe Photoshop CS6 for visualization purposes only and were applied uniformly across image panels.

A Zeiss LSM800 laser scanning confocal microscope was used for fluorescent image acquisition by using 10 x (Plan-Apochromat10 × /0.45), 20 x (Plan-Apochromat 20 × /0.8), and 40 x (EC Plan-Neofluar 40× / NA0.75) objectives. Images were then adjusted using the Zen Blue Edition software.

### Cell density evaluation of c-Fos positive neurons

c-Fos immunolabeled sections obtained from the pup-induced neuronal activation experiments were used to calculate activated cell densities. Microscopic images were analyzed using the open-source bioimage analysis software QuPath v0.4.3 software (University of Edinburgh and Queen’s University Belfast) (Bankhead et al., 2017). The cell detection tool was applied to identify c-Fos+ cell nuclei, while the cell counting tool was used for the quantification of fluorescently double-labeled (c-Fos/Calbindin, c-Fos/PTH2) cells.

### Fiber density measurements of anterograde tract-tracing

For quantitative analysis of fiber density, fluorescent mCherry immunopositivity was evaluated using QuPath v0.4.3. The machine learning–based pixel classification tool was applied to automatically quantify the area density of the anterogradely labeled mCherry+ nerve fibers originating from LSv^Cb+^ neurons across the whole mouse brain. Fiber density was calculated as the proportion of mCherry-immunolabeled fiber area relative to the total analyzed area within each region. Then, the relative contribution of each brain region to the total anterogradely labeled fiber area was calculated and expressed as a percentage.

### Correlative light and electron microscopy

We performed correlative light and electron microscopy to confirm the synaptic connection between PTH2+ terminals and GABAergic neurons by using VGAT-Cre-ZsGreen mouse brain. Mice brains were perfused with a solution containing 4% paraformaldehyde, 0.05% glutaraldehyde, and 0.2% picric acid in PB. 50 µm thin vibratome sectioned brain slices were washed, then cryoprotected overnight first in 15% sucrose, then in 30% sucrose PB solution.

The sections were freeze-thawed over liquid nitrogen for epitope retrieval. 1% sodium-borohydride, 50 mM ammonium-chloride, and 50 mM glycine solution was applied followed by blocking in a solution containing 10% FBS, 5% BSA in PB, and incubating in PTH2 antibody at a dilution of 1:1000 for 48 hours. After washing with PB containing saline, biotin-conjugated secondary antibody (1:500, Jackson Immunoresearch), then avidin-biotin complex (ABC) solution (1:500, Vector Laboratories) was used. Developing happened using A568-conjugated tyramide solution (1:5000) in the presence of 0.001% hydrogen peroxide, and then neurons showing ZsGreen expression and closely surrounded by PTH2+ signal were identified by AxioImager Z1 (CarlZeiss) fluorescent microscope equipped with ApoTome1 unit. After fluorescent image acquisition, the sections were subjected to a peroxidase-based signal developing step. Therefore, a second ABC reagent solution was used for 6 hours. Then, a peroxidase-based reaction by applying NiDAB (DAB peroxidase substrate kit, SK-4100, Vector Laboratories) was performed to develop PTH2-related signal appropriate for the detection with electron microscope.

Brain slices were post-fixed in 1% osmium-tetroxide water solution. Following dehydration steps, sections were embedded in Durcupan^TM^ ACM (Sigma-Aldrich) resin. We identified previously selected GABAergic neurons by light microscope. After re-embedding the area of interest, it was sectioned at 70 nm with a Reichert-Jung ultramicrotome. Sections were collected on formvar film-coated copper slot grids and contrasted with lead citrate.

### Electron microscopic image acquisition

Electron microscopic images were acquired by a Jeol Jem 1011 transmission electron microscope equipped with a Morada camera. The electron microscope was operating at 80 kV. Image adjustments such as brightness and contrast were applied using Adobe Photoshop CS6.

### Quantification and statistical analysis

Statistical analysis of the histological data was conducted with RStudio (version 2025.09.0 Build 387) with the packages lme4, lmerTest, and emmeans. Linear mixed-ANOVA (LM) was used to assess examination groups (mother mice in response to pups vs. mother mice in the absence of pups) and cell type effects on cell densities. The models included fixed effects for the experimental group and the type of immunolabeled cells. The animal ID was included as a random intercept to account for interindividual variability. Post hoc comparisons of estimated marginal means were then applied to test group differences. Normality and homogeneity of variance of model residuals were assessed using the visual inspection of Q–Q plots and residuals versus fitted values plots. Statistical analyses for data collected during 30-minute video recordings of spontaneous maternal behavior tests were assessed by using Two-way ANOVA with Tukey’s multiple comparison test. For other behavioral tests, Two-way ANOVA with Sidak’s multiple comparison test (EPM, FST, OF) or unpaired two-tailed Student’s t-tests (total distance travelled in OF) were conducted. Statistical significance was set at p < 0.05 for all statistical analyses. Diagrams were created in GraphPad Prism (v8.0.2).

**Table 1.** Antibodies and histological reagents used in the study.

| Product name | Host species | Catalogue number | Dilution |
| --- | --- | --- | --- |
| **Primary antibodies** |  |  |  |
| Anti-mCherry Antibody | Chicken | Cat. #: CPCA-mCherry (EnCore Biotechnology Inc.)<br>RRID: AB_2572308 | 1:3000 (fluorescent staining and immunoperoxidase reaction) |
| Anti-Cholera Toxin B subunit | Goat | Cat. #: 703 (List Biological Laboratories)<br>RRID: AB_10013220 | 1:10000 (fluorescent staining) |
| Anti-Calbindin | Mouse | Cat. #: 214 011 (Synaptic Systems)<br>AB_2068201 | 1:2000 (fluorescent staining) |
| Anti-Calbindin | Rabbit | Cat. #: CB38 (Swant)<br>RRID: AB_10000340 | 1:3000 (fluorescent staining) |
| c-Fos Antibody | Rabbit | Cat. #: ab190289 (Abcam)<br>RRID: AB_2737414 | 1:5000 (fluorescent staining and immunoperoxidase reaction) |
| c-Fos Antibody | Guinea Pig | Cat. #: 226 004 (Synaptic Systems)<br>AB_2619946 | 1:15000 (fluorescent) |
| anti-PTH2 | Rabbit | AB_1235466 | 1:3000 (fluorescent staining) |
| anti-PTH2R | Rabbit | Gift from Ted B. Usdin | 1:5000 (fluorescent staining and immunoperoxidase reaction) |
| **Secondary antibodies** |  |  |  |
| Alexa Fluor 488<br>AffiniPure Donkey<br>Anti-Rabbit IgG | Donkey | Cat. #: 715-545-152<br>(Jackson ImmunoResearch)<br>AB_2313584 | 1:500 |
| Alexa Fluor 488<br>AffiniPure Donkey<br>Anti-goat IgG | Donkey | Cat. #: 705-545-147<br>(Jackson ImmunoResearch)<br>RRID: AB_2336933 | 1:500 |
| DyLight 549 Anti-Mouse | Horse | Cat. #: DI-2549-1.5 (Vector<br>Laboratories Inc.) | 1:500 |
| Alexa Fluor 594<br>AffiniPure Donkey<br>Anti-Rabbit IgG | Donkey | Cat. #: 711-585-152<br>(Jackson ImmunoResearch)<br>RRID: AB_2340621 | 1:500 |
| Alexa Fluor 594<br>AffiniPure Donkey<br>Anti-Chicken IgY | Donkey | Cat. #: 703-585-155<br>(Jackson ImmunoResearch)<br>RRID: AB_2340377 | 1:500 |
| Alexa Fluor 594<br>AffiniPure Donkey<br>Anti-Guinea pig IgG | Donkey | Cat. #: 706-585-148<br>(Jackson ImmunoResearch)<br>RRID: AB_2340474 | 1:500 |
| Alexa Fluor 647<br>AffiniPure Donkey<br>Anti-Guinea pig IgG | Donkey | Cat. #: 706-606-148<br>(Jackson ImmunoResearch)<br>AB_2340477 | 1:500 |
| Biotin-SP (long<br>spacer) AffiniPure<br>Donkey Anti-Chicken IgY (IgG) | Donkey | Cat. #: 703-065-155<br>(Jackson ImmunoResearch)<br>RRID: AB_2313596 | 1:1000 |
| Biotin-SP (long<br>spacer) AffiniPure<br>Donkey Anti-Rabbit<br>IgY (IgG) | Donkey | Cat. #: 711-065-152<br>(Jackson ImmunoResearch)<br>RRID: AB_2340593 | 1:1000 |
| **Solutions** |  |  |  |
| DAPI | - | D9542 (Merck, KGaA, Darmstadt, Germany) | 1:8000 for fluorescent staining |
| VECTASTAIN® Elite ABC-HRP Kit, Peroxidase | - | PK-6100 (Vector Laboratories, Burlingame, CA, USA) | 1:500 (immunoperoxidase reaction) |
| Aqua-Poly/Mount | - | 18606 (Polysciences Inc.) | - |
| DePeX mounting medium | - | 06522 (Sigma Aldrich) | - |

## Results

### Neurons in the ventral subdivision of the lateral septum are activated during pup interaction

For mapping the neuronal activation pattern following pup-exposure in mice, we used c-Fos as a neuronal activation marker. To define pup-induced activated neurons, mouse dams were deprived of their pups for 22 hours to reduce c-Fos expression then pups were returned to the homecage for 2 hours. Control dams were not reunited with their pups to determine the baseline activation level (Fig. 1. A). We examined brain areas previously suggested or described to be involved in maternal behavior regulation such as the prefrontal cortex (PFC), the ventral subdivision of lateral septum (LSv), the medial preoptic area (MPOA), the paraventricular hypothalamic nucleus (PVN), the medial amygdaloid nucleus (MeA), the posterior intralaminar thalamic nucleus (PIL) and the ventrolateral subdivision of the periaqueductal grey (vlPAG). Mixed-effects ANOVA revealed a significant main effect of the experimental group (*F*(1, 8) = 120.07, *p* < 0.001), the brain area (*F*(6, 116) = 62,23, *p* < 0.001), as well as the interaction between the experimental groups and brain areas (*F*(6, 116) = 39.902, *p* < 0.001) was significant. The post-hoc analysis revealed that all investigated brain areas showed significantly increased density of c-Fos+ neurons following pup-exposure compared to pup deprivation (in the case of LSv, MPOA, PVN and PIL: \*\*\**p* < 0.0001, vlPAG: *p* = 0.0011, PFC: *p* = 0.0344, MeA: *p* = 0.0147). Among them, one of the lesser investigated brain areas regarding maternal regulation is the LSv. However, based on the c-Fos study, LSv is the third most activated among these brain areas and shows a similar neuronal activation level as the MPOA and PVN, both central regulators of maternal care, suggesting the involvement of the LSv in maternal control.

**Fig. 1:**
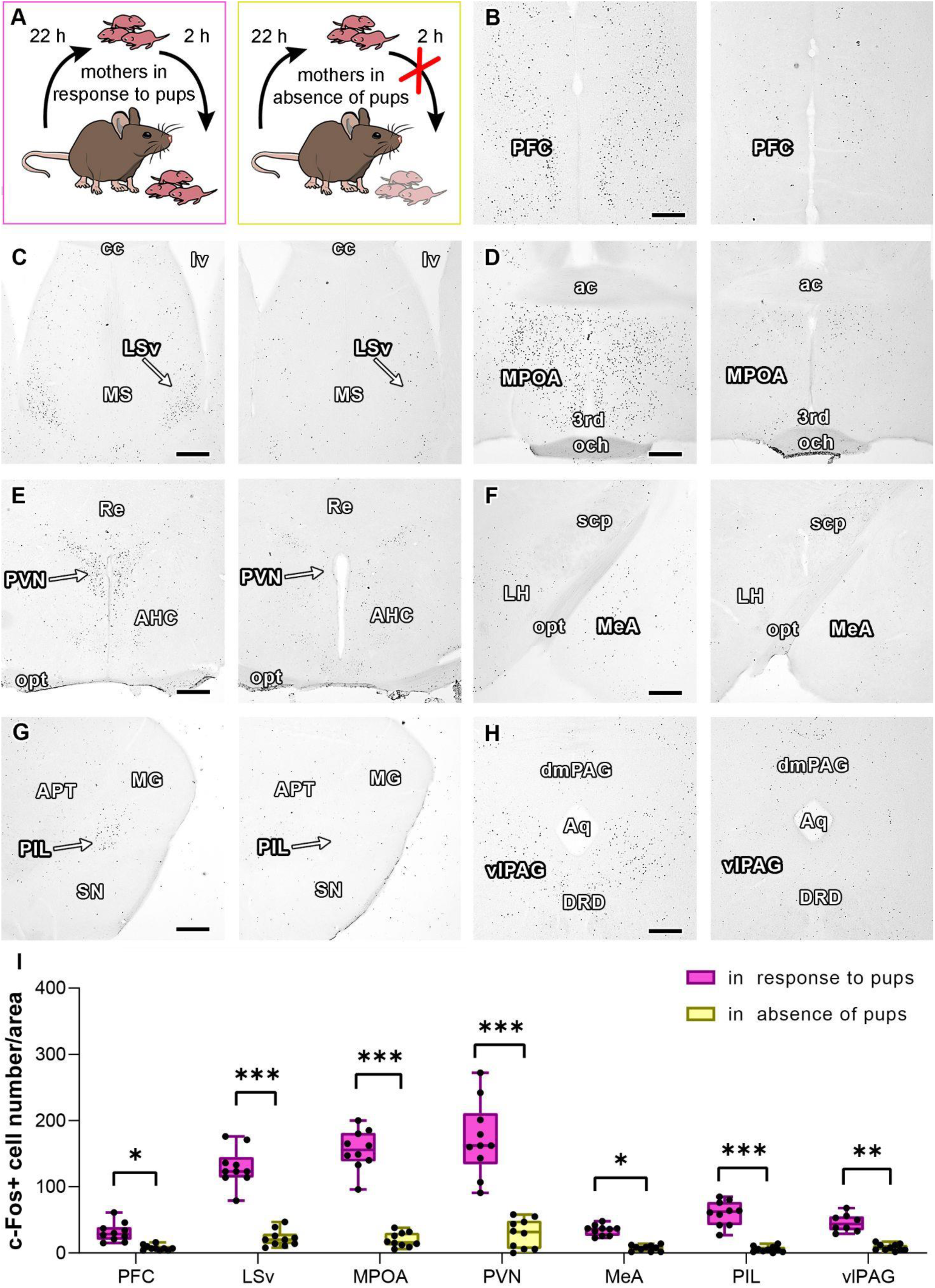
Distribution of the pup-induced c-Fos activated neurons in mother mice during pup-exposure and pup-deprivation. **A:** Graphic illustration of the experimental protocol in pup-exposed mice dams and pup-deprived control ones. **B-H:** Representative microscopic images from the examined brain areas in pup-exposed (left) and control animals (right). **I:** Significantly elevated c-Fos activation was seen in response to pups compared to full deprivation in the illustrated brain areas and nuclei (LSv, MPOA, PVN, and PIL: \*\*\**p* < 0.0001, vlPAG: *p* = 0.0011, PFC: *p* = 0.0344, MeA: *p* = 0.0147, mixed-effects ANOVA with post hoc analysis using estimated marginal means, n = 6 per group). Box plots display whiskers extending to the minimum and maximum observed values. The horizontal lines above bars indicate that p value is significant (*: *p* < 0.05, **: 0.001 < *p* < 0.01, ***: *p* < 0.001). Scales: 500 μm. Abbreviations: 3rd: third ventricle, ac: anterior commissure, AHC: anterior hypothalamic nucleus central part, APT: anterior pretectal nucleus, Aq: aqueduct, cc: corpus callosum, DRD: dorsal part of the dorsal raphe nucleus, LH: lateral hypothalamus, LSv: ventral subdivision of the lateral septal nucleus, lv: lateral ventricle, MeA: medial amygdaloid nucleus, MG: medial geniculate nucleus, MPOA: medial preoptic area, MS: medial septal nucleus, och: optic chiasm, opt: optic tract, dmPAG: dorsomedial periaqueductal grey, vlPAG: ventrolateral periaqueductal grey, PFC: prefrontal cortex, PIL: posterior intralaminar thalamic nucleus, PVN: paraventricular nucleus of the hypothalamus, Re: reuniens thalamic nucleus, scp: superior cerebellar peduncle, SN: substantia nigra.

### Pup-induced c-Fos activated LSv neurons are calbindin-positive

The group of c-Fos+ neurons following pup interaction found in the LSv showed a similar localization pattern as the previously described calbindin+ (Cb^+^) neuron population in the area (Zhao et al., 2013). Therefore, we hypothesized that Cb^+^ neurons exhibit pup-induced c-Fos activation in the LSv. In order to describe the activation pattern of this Cb^+^ neuron population, we used the aforementioned pup-deprivation protocol (Fig. 2. A, C), and then double labelled the slices with c-Fos and calbindin antibodies as seen on the representative microscopic images (Fig. 2 B, D). We revealed differences between the examined groups (*F*(1, 10) = 28.887, *p* < 0.001) and cell types (*F*(1, 10) = 10.646, *p* = 0.0085) by using a linear mixed-effect model. The analysis of the fluorescently labeled samples also confirmed the previously described increase of activated neuronal density in the LSv (p < 0.001). Moreover, we found that the density of activated Cb^+^ neurons was also significantly higher in response to the pups compared to complete pup deprivation (Fig. 2. E, *p* = 0.0015), although a larger difference was observed in c-Fos+ cell density, based on the interaction effect between the examined groups and observed cell types (*F*(1, 10) = 10.646, *p* = 0.0085).

**Fig. 2:**
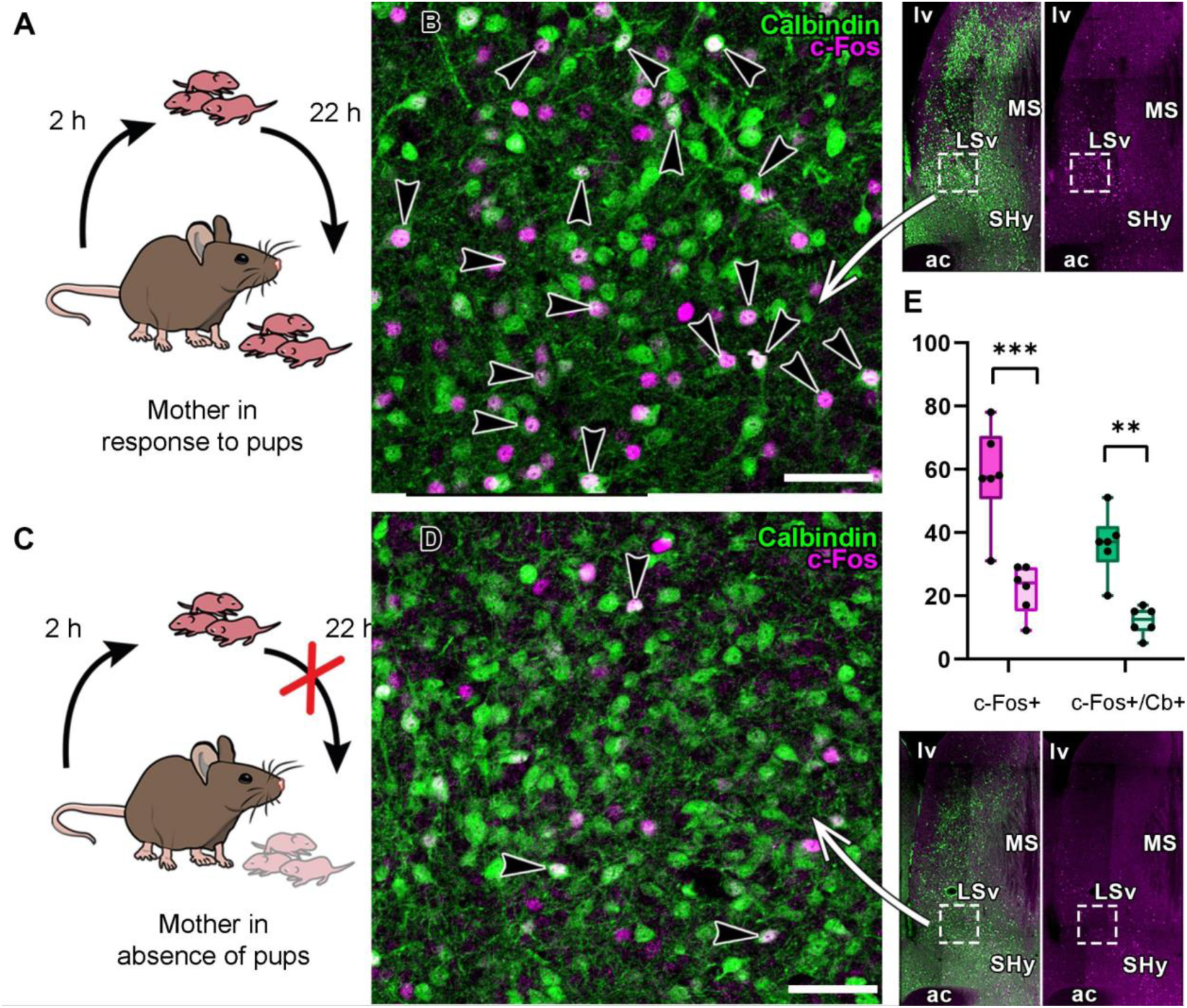
Calbindin-positive neuron population of the ventral subdivision of the lateral septum exhibits pup-induced c-Fos activation in mouse dams. **A,C:** Mother mice were deprived of their pups for a day, then were either pup-exposed (**A**) or remained pup-deprived (**C**) for 2 hours before sacrifice. **B, D:** Representative confocal microscopic images from the two experimental groups indicate elevated c-Fos activation in the septal calbindin neuron population in pup-exposed dams (B) compared to pup-deprived ones (D). Right inlet panels show the low magnification images of the LS indicating the area used for higher magnification images appearing on the left panels. **E:** The density of c-Fos+ and double labeled (Cb+/c-Fos+) cells in the examined areas are shown on the graph. Area = 0.1 mm2. Box plots display whiskers extending to the minimum and maximum observed values. The horizontal lines above bars indicate that p value is significant (*: *p* < 0.05, **: 0.001 < *p* < 0.01, ***: *p* < 0.001). \*\*\**p* < 0.001, 0.01 < \**p* < 0.05. Scales: B, D: 50 μm. Abbreviations: ac: anterior commissure, MS: medial septum, lv: lateral ventricle, LSv: ventral subdivision of the lateral septum, SHy: septohypothalamic nucleus.

The somatic area of both the pup-induced activated and non-activated LSv^Cb+^ neurons was quantified to analyze morphological differences in cell size within the Cb neuronal populations. The unpaired t-test did not reveal a significant difference (*p* = 0.1298, n = 20 per cell type) between the somatic area of c-Fos-positive (area: 128.37 ± 27.04 µm^2^ (SD)) and c-Fos-negative (area: 117.30 ± 17.05 µm^2^ (SD)) neurons.

### Calbindin-positive neurons of the lateral septum have distinguished electrophysiological properties compared to calbindin-negative septal neurons

Since LSv^Cb+^ neurons are highly activated in response to pups, we addressed whether these neurons have any specific characteristic that can distinguish them from the other septal neuron populations (Fig. 3. A). To gain some morphological information about these neurons, we filled them with biocytin and visualized them fluorescently. LSv^Cb+^ neurons exhibit smaller and elliptically shaped cell bodies and less characteristic dendritic arborization than LSv^Cb-^neurons. The somatic area of the biocytin-filled LSv^Cb+^ neurons was similar (somatic area: 119.20 μm ± 14.89 μm SD, n = 3) to that of maternally activated cells (Fig. 3. B-C). Electrophysiological characterization of calbindin-positive (Cb^+^) and -negative (Cb^-^) neurons in the LSv revealed that both neuronal populations display regular firing in response to rectangular current steps (Fig. 3. D, E). There was no apparent difference in spike shape (Fig. 3. G, I), however, in the Cb^+^ population, the mean amplitude and rise slope of spikes was significantly lower than in Cb^-^ neurons (Fig. 3. L and M). In some neurons, a moderate voltage sag was found at negative current steps (Fig. 3. E), while in other neurons, this was absent (Fig. 3. D), however, there was no systematic difference between the two neuronal populations in this parameter (Fig. 3. K). In contrast, the membrane resistance of Cb^+^ neurons was significantly higher than the resistance in the Cb^-^ population (Fig. 3. J).

**Fig. 3:**
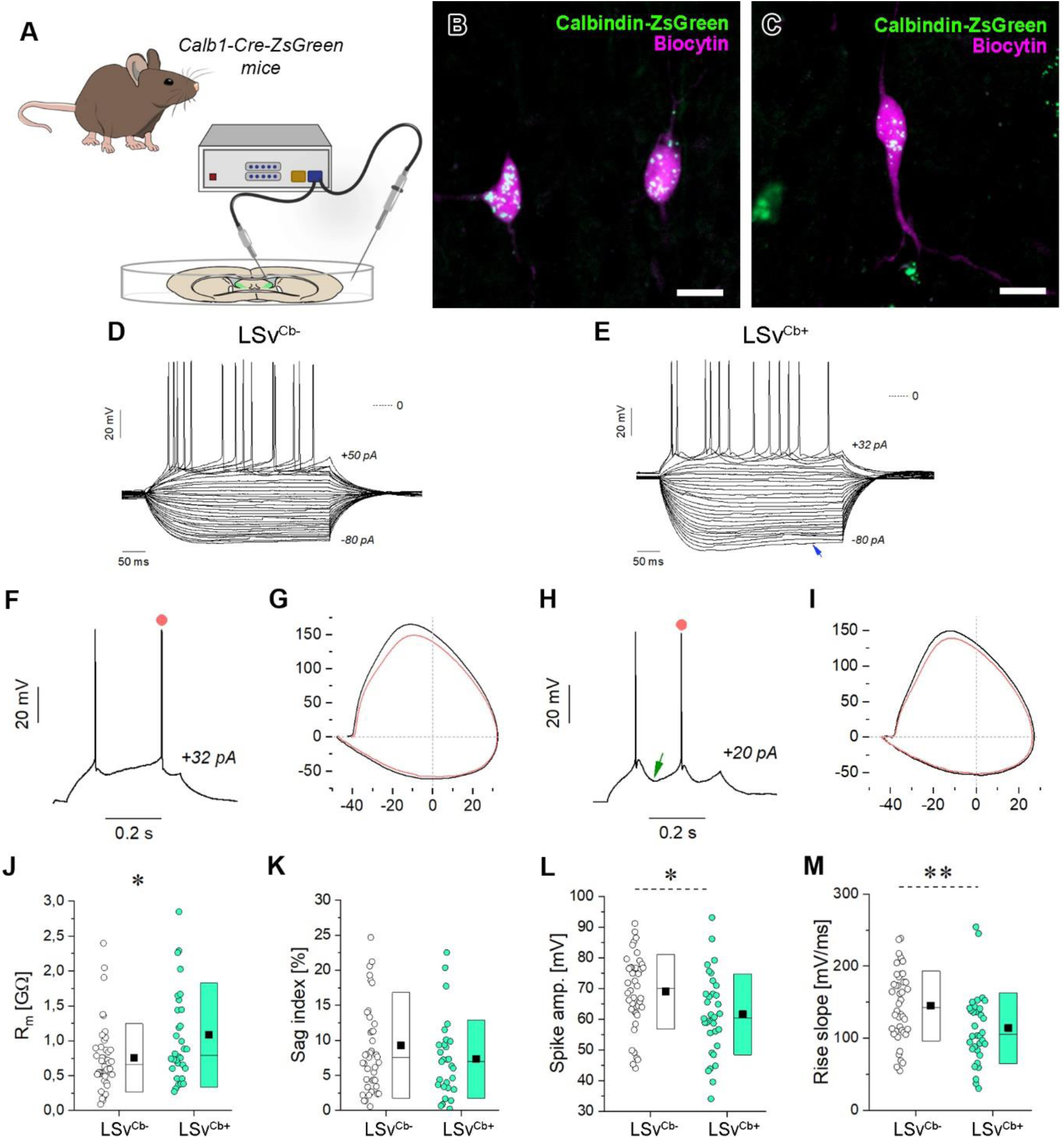
Lateral septum neurons exhibit diverse electrophysiological properties as revealed by their voltage responses under current step stimulation. A: Schematic visualization of the whole-cell patch clamp measurement. **B, C:** We filled calbindin positive neurons with biocytin to receive morphological information about them. Representative confocal microscopic images (**B, C**) of these neurons show smaller elliptically shaped cell bodies with less defined dendritic arborization. **D-E:** Incrementing levels of rectangular current steps applied in whole-cell conditions elicited the voltage traces of calbindin negative (LSv^Cb-^) and positive (LSv^Cb+^) cells, respectively. Selected suprathreshold voltage traces in **F** and **H** demonstrate the action potentials of both cell types. The LSv^Cb+^ neuron exhibits moderate voltage sag (blue arrow) under negative current steps (**E**) and pronounced spike afterhyperpolarization (**H**, green arrow). Phase portraits of the action potentials of **F** and **H** are shown in **G** and **I**, respectively (red trajectories for the second spikes). **J-M:** Four physiological parameters extracted from the voltage responses are compared for the LSv^Cb-^ and LSv^Cb+^ neurons. The membrane resistance of the LSv^Cb+^ cells appears significantly higher than that of the LSv^Cb-^ neurons (**J**). Voltage sag index (**K**), indicative of the strength of the h-current in the neurons ranges from 0 to 25% but is not significantly different for the two populations. Spike amplitude in L and the upshoot slope of the action potentials (**M**) differ slightly in LSv^Cb-^ and LSv^Cb+^ neurons. (*: *p* < 0.05, **: *p* < 0.01).

### LSv^Cb+^ neurons promotes pup-licking behavior in female mice

Since LSv^Cb+^ neurons exhibit prominent activation in the wake of interaction with pups and have some unique electrophysiological characteristics compared to LSv^Cb-^neurons, we addressed the functional role of these neurons in the regulation of maternal behavior. We chemogenetically inhibited the Cb+ neurons of the LSv. For this, we injected an inhibitory DREADD (Designer Receptors Exclusively Activated by Designer Drugs) and fluorescent protein (mCherry) encoding AAV expressing in a Cre dependent manner into the LSv of Calbindin-Cre female mice (Fig. 4. A, B). The control group received a similar viral vector without the gene of the inhibitory receptor. To verify the viral efficiency, slices were double immunolabelled with calbindin and mCherry as seen in the representative images (Fig. 4. C). Both examined groups were prepared from virgin females sensitized beforehand with pups. Test schedule was determined in a way to ensure self-control among the examined animals to reduce the individual effects of behavior meaning that all animals had two control days (day 1 and day 5) on which they received only the vehicle and one test day (day 3) receiving the exogenous ligand (clozapine-N-oxide, CNO) injection (Fig. 4. D). Analyses followed by statistical comparison of control and test days were made in each case. Maternal care was examined using spontaneous maternal behavior test, where mice could freely interact with the pups, and then pup-related behavior elements were examined (Fig. 4. D). Inhibition of LSv^Cb+^ neurons decreased the time spent with licking of pups compared to control days (Fig. 4. G-H, *p* = 0.0066). No significant changes were found in other elements (Fig. 4. E-F; pup grooming: *p* = 0.9705; pup sniffing: *p* = 0.3576; nest building: *p* = 0.7734) or in the control groups (Fig. 4. G-H; pup grooming: *p* = 0.7955; pup licking: *p* = 0.2781; pup sniffing: *p* = 0.9588; nest building: *p* = 0.8774). To verify that the observed change is not related to other behavioral alterations, supplementary tests of anxiety- and depression-like behaviors were completed, such as open field, elevated plus maze, and forced swim tests (Fig. 4. I). No changes in the time spent in the corner (*p* > 0.9999), wall (*p* = 0.9549) and middle region (*p* = 0.9667) of the open field apparatus and in the travelled distance (*p* = 0.0885) were found comparing inhibited and control animals (Fig. 4. J). Similarly, both inhibited and control female mice spent similar amount of time in the open (*p* = 0.9867) and closed arms (*p* = 0.4188) and center region (*p* = 0.9993) of the elevated plus maze (Fig. 4. K). Regarding depression-like behavior, mice with inhibited LSv^Cb+^ neurons showed similar behavior as the control mice in forced swim test (Fig. 4. L; swimming: *p* = 0.4154; climbing: *p* = 0.9998; floating: *p* = 0.4623). These additional tests strengthened the specific results of the spontaneous maternal behavior test, as the observed significant change was not caused indirectly by other behavioral changes.

**Fig. 4:**
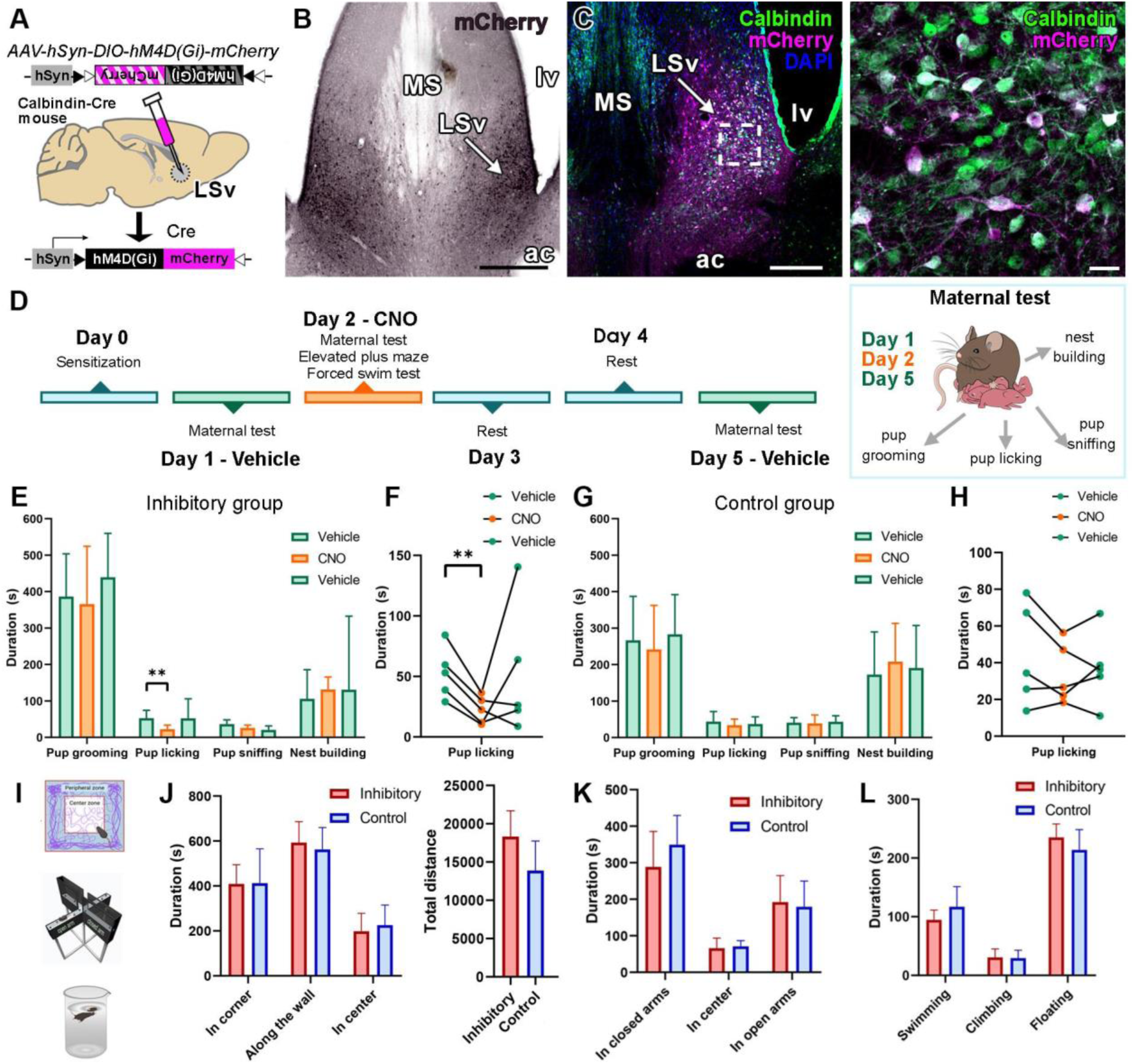
Inhibition of calbindin neurons in the ventral part of the lateral septum reduces pup licking behavior in female mice. **A, B:** Schematic illustration describing the viral chemogenetic method (**A**) and a representative image of the injection site in the ventral subdivision of the lateral septum (LSv) (**B**). Scale: 500 μm. Abbreviations: ac: anterior commissure, lv: lateral ventricle, MS: medial septal nucleus. **C:** Confocal microscopic images verify the proper operation of the virus construct. Scales: C left panel: 500 μm, right panel: 20 μm. **D:** Visualization of the used behavioral test schedule and the examined spontaneous maternal behavior test. **E-H:** Effect of inhibiting LSv^Cb+^ neurons on the time spent with spontaneous maternal behavior elements. Inhibition significantly decreased the duration of pup licking behavior (Two-way ANOVA with Tukey’s multiple comparison test, **: *p* = 0.0066, n = 5) on CNO administration day compared to vehicle days (**E-F**). This shift in duration cannot be seen in the non-inhibited control female group (**G-H**, n = 5). **I:** Graphic illustration of the used anxiety- and depression-like behavior test, as open field, elevated plus maze, and forced swim test. **J:** Results of open field test show no difference in time spent in the different areas of the open field apparatus and in total travelled distance between inhibited and control females. **K:** Inhibited animals spent the same amount of time in the open and closed arms of the apparatus as control ones. **L:** During forced swim test, no difference in behavior was observed comparing the two experimental groups.

### LSv^Cb+^ neurons project to brain areas involved in maternal care

Behaviors are usually regulated by complex neural networks. Therefore, we examined the neuronal connections of LSv^Cb+^ neurons. Injection of a viral vector Cre dependently expressing mCherry fluorescent protein into the LSv of Calbindin-Cre female mice revealed output areas of LSv^Cb+^ neurons (Fig. 5.). After the quantitative analysis, the medial preoptic area (MPOA), a center for maternal behavior regulation, was confirmed to be the strongest output area (12.7%) followed by the anterior commissural nucleus (AC, 7.7%), the central part of the anterior hypothalamic area (AHC, 7.5%), medial tuberal nucleus (MTu, 7.2%), supraoptic nucleus (SO, 5.7%), ventrolateral preoptic nucleus (VLPO, 5.6%), and terete hypothalamic nucleus (Te, 5.1%). Further prominent projection sites were determined: the suprachiasmatic nucleus (SCh, 4.6%), ventral hippocampus (vHipp, 4.6%), lateral hypothalamic area (LH, 3.5%), lateroanterior hypothalamic nucleus (LA, 3.3%), anterior part of the anterior hypothalamic area (AHA, 3.2%), tuber cinereum area (TC, 3.1%), bed nucleus of stria terminalis (BNST, 3.2%), dorsal tuberomamillary nucleus (DTM, 3.0%), ventral premamillary nucleus (PMV, 3.0%), arcuate nucleus (Arc, 2.9%), and dorsomedial hypothalamic nucleus (DMH, 2.8%) in decreasing order. A lower density of fibers was found in the supramamillary nucleus (SuM, 2.4%), the posterior part of the anterior hypothalamic area (AHP, 2.4%), the subparaventricular zone of the hypothalamus (SPa, 2.3%), the posterodorsal part of the medial amygdaloid nucleus (MePD, 1.7%), posterior hypothalamic nucleus (PH, 1.4%), the anterodorsal part of the medial amygdaloid nucleus (MeAD, 0.6%), periaqueductal grey (PAG, 0.4%), the medial parvicellular part of the paraventricular hypothalamic nucleus (PaMP, 0.4%), and the interfascicular nucleus (IF, 0.4%).

**Fig. 5:**
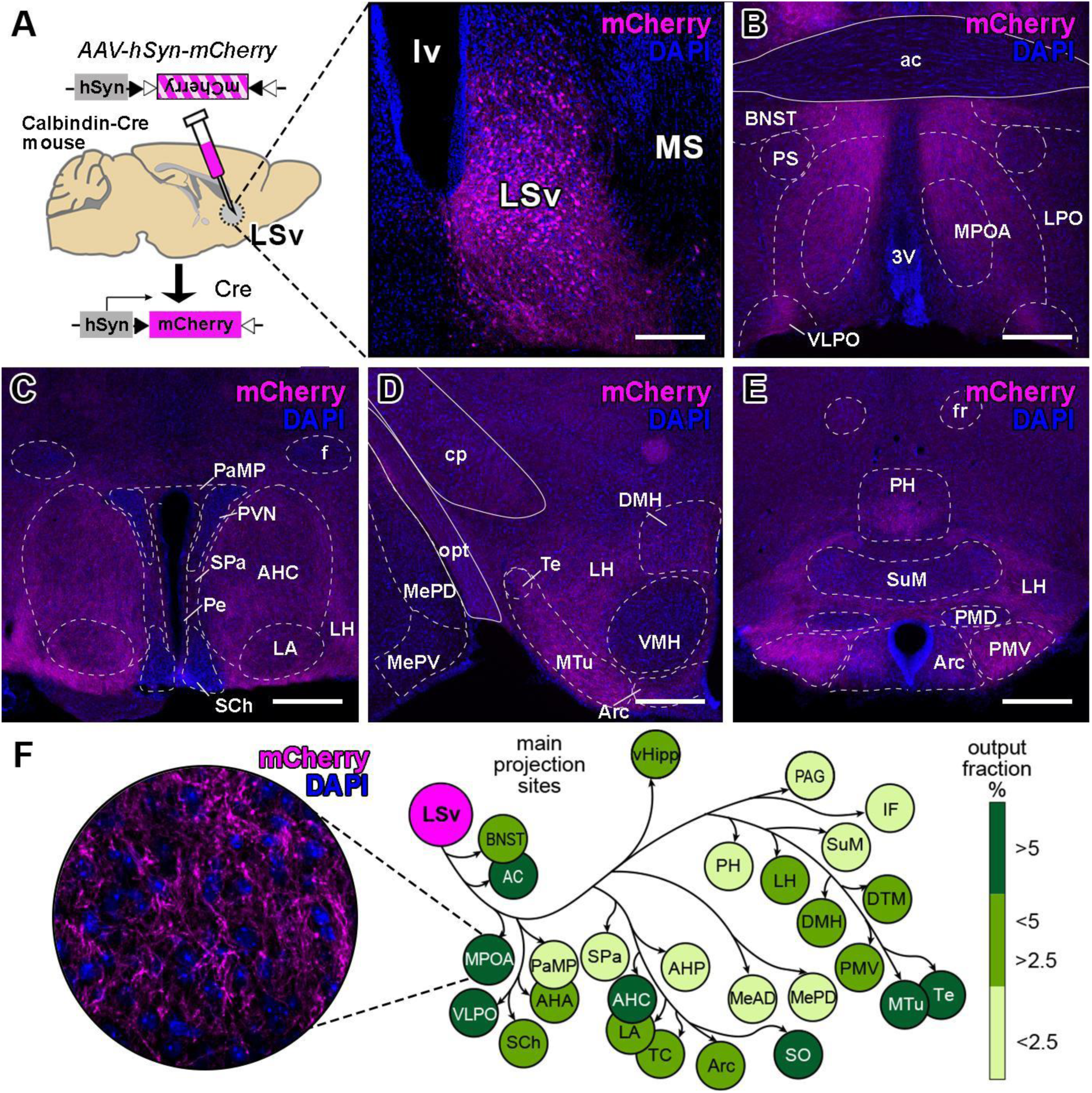
Mapping of the output areas of LSv^Cb+^ neurons. **A:** Visualization of the anterograde tracing from LSv^Cb+^ neurons with a representative image of the injection site. Scale: A: 200 μm for inlet. **B-E:** Representative images of output areas containing mCherry-positive fibers (magenta channel). Scale: B-E: 200 μm. **F:** Overview of the outputs of LSv^Cb+^ neurons in virgin female mice (n=2). Fractions are shown in mean percentages and are color-coded with the darkest green indicating the strongest output. The inlet shows a higher magnification image from the densest output area, the medial preoptic area (MPOA, 12.7%). Scale: F: 20 μm for inlet. Abbreviations: 3V: third ventricle, ac: anterior commissure, AC: anterior commissural nucleus, AHA: anterior part of the anterior hypothalamic area, AHC: central part of the anterior hypothalamic area, AHP: posterior part of the anterior hypothalamic area, Arc: arcuate nucleus, BNST: bed nucleus of stria terminalis, cp: cerebral peduncle, DMH: dorsomedial hypothalamic nucleus, DTM: dorsal tuberomamillary nucleus, fr: fasciculus retroflexus, vHipp: ventral hippocampus, IF: interfascicular nucleus, LA: lateroanterior hypothalamic nucleus, LH: lateral hypothalamic area, LSv: ventral subdivision of the lateral septum, LPO: lateral preoptic nucleus, lv: lateral ventricle, MeAD: anterodorsal part of the medial amygdaloid nucleus, MePD: posterodorsal part of the medial amygdaloid nucleus, MePV: posteroventral part of the medial amygdaloid nucleus, MPOA: medial preoptic area, MS: medial septum, MTu: medial tuberal nucleus, opt: olivary pretectal nucleus, PAG: periaqueductal grey, PaMP: medial parvicellular part of the paraventricular hypothalamic nucleus, Pe: periventricular hypothalamic nucleus, PH: posterior hypothalamic nucleus, PMD: dorsal premamillary nucleus, PMV: ventral premamillary nucleus, PS: parastriatal nucleus, SCh: suprachiasmatic nucleus, SPa: subparaventricular zone of the hypothalamus, SO: supraoptic nucleus, SuM: supramamillary nucleus, TC: tuber cinereum area, Te: terete hypothalamic nucleus, VMH: ventromedial hypothalamic nucleus, VLPO: ventrolateral preoptic nucleus.

### Anatomical examination of the LSv^Cb+^-MPOA pathway

As the major output region of LSv^Cb+^ neurons is the MPOA, an area critical to maternal behavior, we further examined this connection for its role in licking behavior regulation. A retrogradely transported neuronal tracer was injected into the MPOA of female mice (Fig. 6). Then the number of retrogradely labelled Cb^+^ neurons was counted in the LSv. A high proportion of the retrogradely labeled MPOA-projecting LSv neurons (CTB+) showed calbindin positivity (60.9 ± 10.3%). The quantification of the somatic area revealed that MPOA-projecting LSv^Cb+^ neurons were similar in size (the mean of somatic area: 119.30 ± 23.43 μm^2^ (SD)) to maternally activated cells.

**Fig. 6:**
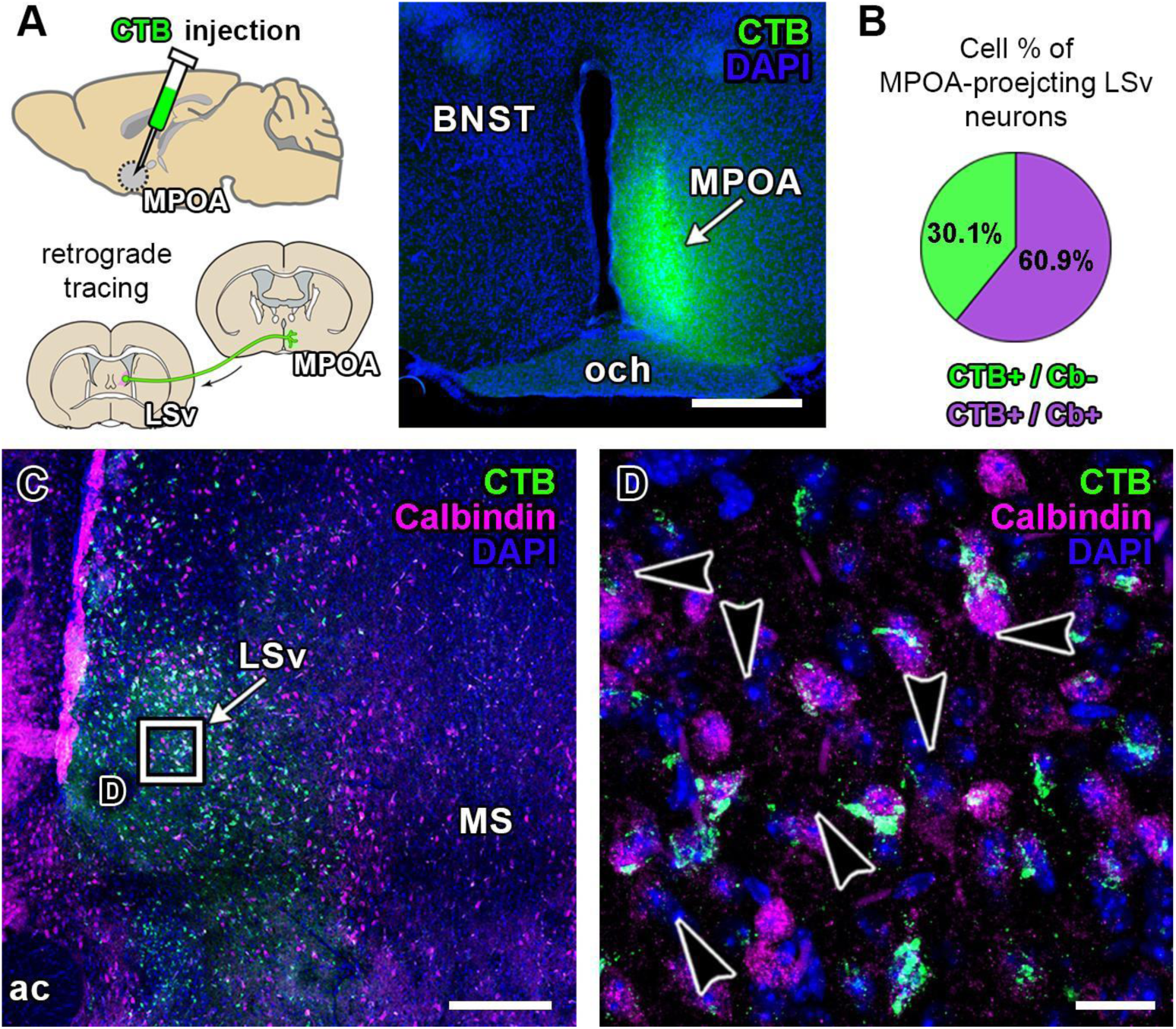
Prominent septal input of the medial preoptic area (MPOA) originates from LSv^Cb+^ neurons. **A:** The experimental design of the retrograde tract tracing shows the injection of cholera toxin beta subunit (CTB) into the MPOA. Scale: 500 µm. Abbreviations: BNST: bed nucleus of the stria terminalis, och: optic chiasm. **B:** The density of the retrogradely labelled (CTB+) neurons was determined based on the presence or absence of calbindin positivity in double immunolabeled sections. Cb^+^ neurons represent one of the major septal inputs to the MPOA, as 60.9 ± 10.3% of the MPOA-projecting cells (CTB+ cells) are Cb positive. **C:** The low magnification confocal image shows the distribution pattern of the retrogradely labeled cells in the LSv. Scale: 250 µm. Abbreviations: ac: anterior commissure, LSv: ventral subdivision of the lateral septal nucleus, MS: medial septal nucleus. **D:** The magnified part of panel C (indicated by the rectangle) demonstrates that a high number of the MPOA-projecting neurons are double-labeled with CTB and Cb (arrowheads). Scale: 20 µm.

### Determining maternally relevant activation pathway of LSv^Cb+^ neurons via PTH2+

Maternally induced PTH2 is involved in the control of maternal behaviors, therefore, we examined whether PTH2+ fibers in the LSv have a connection with pup-induced c-Fos activated Cb^+^ neurons in mother mice (Fig. 7. A-C). We found that 79.4 ± 3.5 % of the activated LSv^Cb+^ neurons are closely apposed by PTH2+ fibers (Fig. 7. B). Furthermore, on average, 15.25±6.4 putative synaptic puncta were found around the cell bodies of c-Fos+/Cb+ LSv neurons (Fig. 7. C, D). In addition, the potential synaptic relationship between pup-induced c-Fos+ LSv^Cb+^ neurons and PTH2+ fibers was confirmed in the rat brain (Fig. 7. D). We investigated the localization of the receptor of PTH2 (PTH2R), as well, and found that PTH2R has a similar expression pattern as PTH2+ fibers in the LSv (Fig. 7. E). Double labeling of PTH2R with calbindin revealed that many LSv^Cb+^ neurons also show PTH2R positivity (Fig. 7. F, G).

**Fig. 7:**
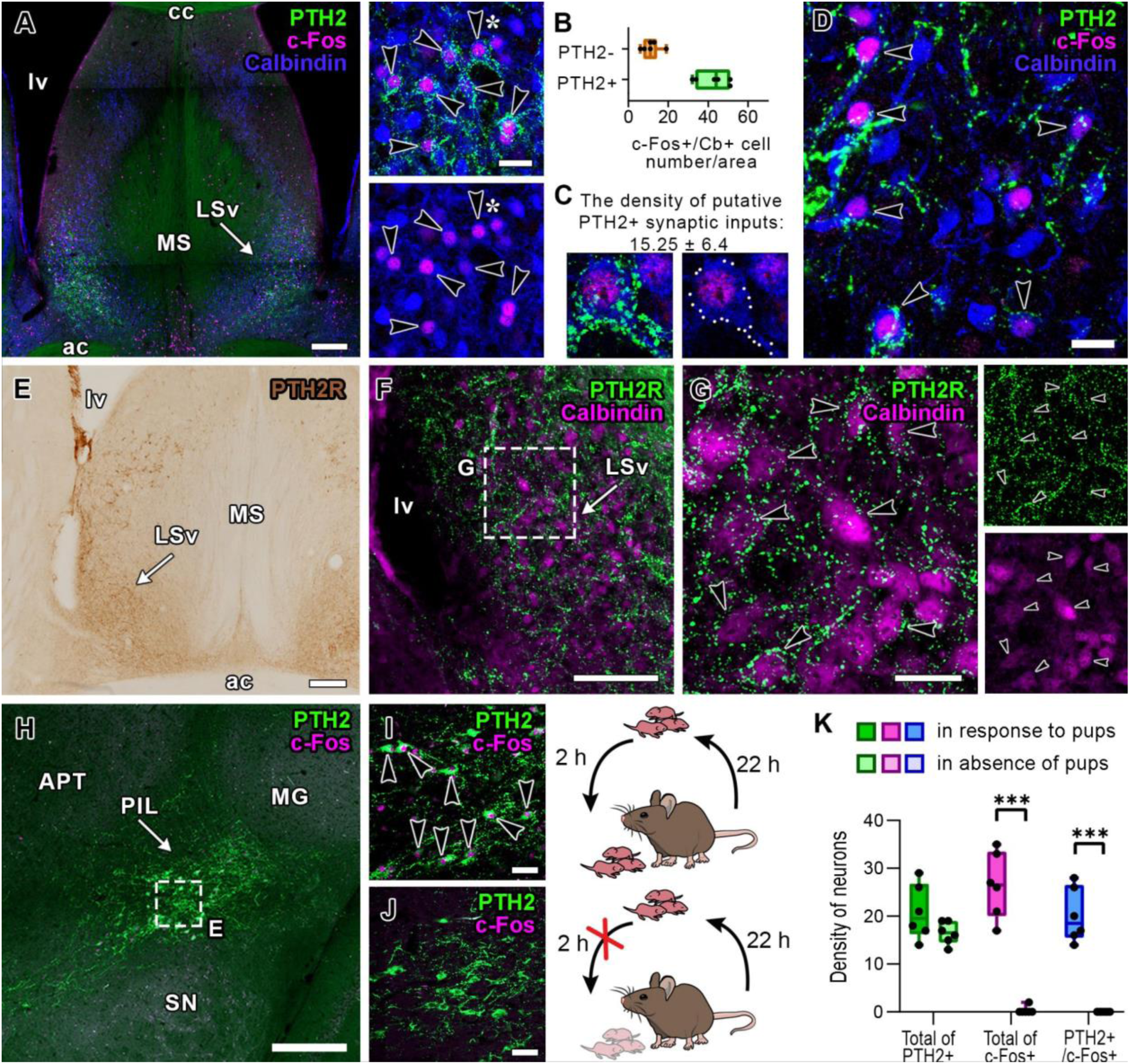
Parathyroid hormone 2 (PTH2)-positive fibers closely appose LSv^Cb+^ neurons activated by pup-exposure. **A:** A low magnification confocal image demonstrates that PTH2+ fibers show overlapping distribution with pup-induced c-Fos activated LSv^Cb+^ neurons, which are closely apposed by PTH2+ fibers. Scale: left panel: 250 µm, right panel: 20 µm. Abbreviations: ac: anterior commissure, cc: corpus callosum, LSv: ventral subdivision of the lateral septum, lv: lateral ventricle, MS: medial septal nucleus. **B:** Pup-induced c-Fos activated LSv^Cb+^ neurons surrounded by PTH2+ fibers exhibited a higher neuronal density (43.0 ± 8.4 SD neurons per mm^2^) compared with neurons lacking PTH2+ fiber apposition (12.0 ± 4.5 SD neurons per mm^2^). **C:** The density of putative synaptic structures showing PTH2 positivity of c-Fos+ LSv^Cb+^ neurons was determined. The representative high magnification image of the same neuron indicated by an asterisk in panel A demonstrates numerous PTH2+ puncta marked by white dots in the right panel. **D:** The same condition can be observed in rat dams, as well, as indicated by the representative confocal microscopic image with black arrowheads pointing at c-Fos+ LSv^Cb+^ neurons surrounded by PTH2+ fibers. **E-G:** Immunolabeling shows the distribution pattern of PTH2 receptor (PTH2R) localization across the LS. The low (**F**) and high (**G**) magnification images of the LSv demonstrate PTH2R immunopositivity of LSv^Cb+^ neurons (pointed by the arrowheads). Scales: E: 250 µm, F: 100 µm, G: 20 µm. **H-J:** Illustrated by schematic graphics, representative confocal microscopic images from the PIL (H) indicate elevated c-Fos activation in mother mice exposed to pups (**I**) compared to pup-deprived ones (**J**). **K:** All PIL^PTH2+^ neurons show c-Fos activation in response to pups in mother mice compared to pup-deprived control mothers (*** *p* < 0.0001, linear mixed-ANOVA with post hoc comparison, n = 6). Box plots display whiskers extending to the minimum and maximum observed values. The horizontal lines above bars indicate that p value is significant (*: *p* < 0.05, **: 0.001 < *p* < 0.01, ***: *p* < 0.001). Scales: 250 µm for the left panel, 20 µm for the right panels. Abbreviations: APT: anterior pretectal nucleus, MG: medial geniculate nucleus, PIL: posterior intralaminar thalamic nucleus, PTH2: parathyroid hormone 2, SN: substantia nigra.

Since we found PTH2+ fibers around the maternally activated LSv neurons, we examined c-Fos-activation in response to pups in PIL^PTH2+^ neurons (Fig. 7. H), especially since we found a significant elevation in the density of c-Fos+ in response to pups in the PIL of mother mice (Fig. 1. G, I).

PIL neurons expressed c-Fos following pup-exposure (Fig. 7. H, I), and this activation terminates when mothers are deprived of their pups (*p* < 0.001) (Fig. 7. J, K). Furthermore, all PIL^PTH2+^ neurons show pup-induced c-Fos-activation and none are activated if the pups are absent (*p* < 0.001) (Fig. 7. K). We revealed a significant effect of the group (*F*(1, 10) = 49.054, *p* < 0.001), cell type (*F*(1,20) = 73.946, *p* < 0.001), as well as their interaction (*F*(1, 20) = 122.097, *p* < 0.001) by using a linear mixed-ANOVA model.

### PTH2+ terminals innervate the maternally involved inhibitory neurons of the LSv

To gain more information about the possible regulatory role of LSv^Cb+^ neurons, we examined what other neurotransmitter is present in these neurons. We fluorescently labelled calbindin on slices from transgenic mouse strain co-expressing ZsGreen fluorescent protein in vesicular GABA transporter (VGAT) containing neurons, which is a marker for GABAergic neurons. Cb+ neurons expressed VGAT suggesting they are part of the inhibitory GABAergic septal neuron population (Fig. 8. A).

**Fig. 8:**
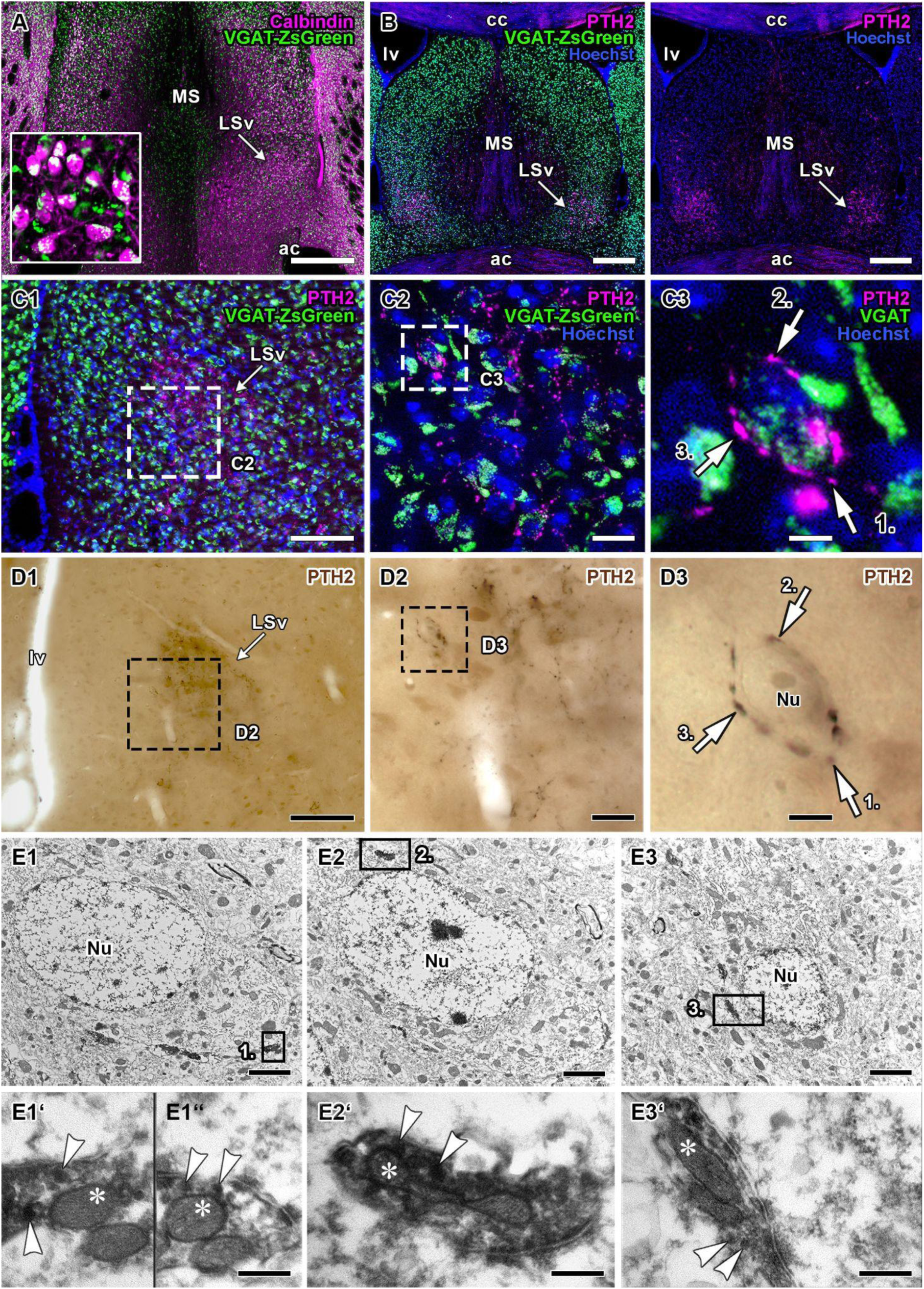
Innervation of GABAergic lateral septal neurons by PTH2+ terminals. **A:** Microscopic image of the LSv of VGAT-ZsGreen female mouse following Cb immunolabelling confirms that LSv^Cb+^ neurons are GABAergic. **B:** Supported by the representative fluorescent microscopy image, GABAergic neurons in the LSv are closely surrounded by PTH2+ fibers suggesting synaptic connection. **C1:** Fluorescent microscopic image of the ventral subdivision of the lateral septum (LSv) following PTH2 immunolabelling of VGAT-ZsGreen mother mouse shows a prominent PTH2+ fiber network located in the area. **C2-3:** Higher magnification images demonstrate that PTH2+ nerve terminals closely appose VGAT+ neuronal cell bodies (marked by numbered white arrows) supposing synaptic contact. **D1:** The same area is shown in panel C1 following immunoperoxidase visualization of PTH2 immunoreaction. **D2:** The panel shows the same area as in panel C2. **D3:** Image of the same neuron as in panel C3. White arrows indicate PTH2+ terminals. **E1-C3:** Low magnification electron microscopic images of serial sections of the neuron are shown in panels C3 and D3. The numbers indicate the same PTH2+ terminals marked in the fluorescent high magnification panel (C3). **E1’-C3’:** High magnification images of the terminals indicated with numbers. Panel E1” is an image of the following ultrathin section of the same synaptic terminal shown in panel E1’. Synaptic dense core vesicles are pointed at by white arrowheads, while mitochondria are signed by asterisks. Scales: C1, D1: 100 μm; C2, D2: 20 μm; C3, D3: 5 μm; E1, E2, E3: 2.5 μm; E1’, E1”, E2’, E3’: 250 nm. Abbreviations: LSv: ventral subdivision of the lateral septum, lv: lateral ventricle, Nu: nucleus.

To determine if PTH2+ fibers found in the area of the GABAergic LSv^Cb+^ neurons (Fig. 8. B) innervate these inhibitory neurons, correlative light and electron microscopy (CLEM) was used on the aforementioned VGAT-ZsGreen transgenic mice strain slices. LSv^VGAT+^ neuron surrounded by PTH2+ terminals was identified first using fluorescent immunohistochemistry of PTH2 (Fig. 7. C1-3). Then, the same neuron was visualized with peroxidase-based immunohistochemistry (Fig. 7. D1-3) and identified using electron microscope (Fig. 7. E1-3) and PTH2+ structures (numbered in the figure) were further examined. Based on high magnification electron microscopic images, the localization of small synaptic and dense core vesicles, puncta adherentia, the synaptic cleft, and the postsynaptic density between the GABAergic neuron and the PTH2+ terminals were demonstrated (Fig. 7 E1’-E3’).

## Discussion

In the current study, we aimed to describe how LSv neurons affect the regulation of maternal behavior and to define their role within the maternal neural circuitry.

### Pup-induced activation in the LSv

The occurrence of c-Fos activation in the LSv in response to reuniting with their litters has been well-documented in rat mothers (Li et al., 1999; Lin et al., 1998; Lonstein et al., 1997) and has also been reported in mice (Hasen & Gammie, 2005; Oláh et al., 2018). Our study confirmed this finding and revealed that c-Fos activation is among the highest in the LSv, suggesting its role in maternal behavior. During reunion, mothers interact with their pups in a stereotyped behavioral pattern that includes increased pup-directed maternal behaviors such as grooming, licking, sniffing, and nursing (Rombaut et al., 2023). However, it is unclear which behaviors and inputs are associated with elevated c-Fos levels in the LSv due to the low temporal resolution of current techniques. Previous studies have suggested that somatosensory inputs from the pups are necessary for the highest level of c-Fos activation in the LSv of rat dams (Lin et al., 1998; Puska et al., 2025). The neurochemical nature of LSv neurons activated by pup exposure remains unknown. Previous studies reported that LSv is enriched in Cb+ neurons (Olucha-Bordonau et al., 2012; Risold & Swanson, 1997a). Here, we demonstrate that the overwhelming majority of pup-induced c-Fos-activated neurons contain calbindin. Using a transgenic approach, we also provided evidence for the GABAergic nature of LSv^Cb+^ neurons. The extant literature on the functional roles of LSv^Cb+^ neurons in biological processes is, as yet, rather sparse. The only literature on this subject that has been published to date mentions the role of these neurons in TrkB signaling (Fawcett et al., 1999) and the influence of these neurons on theta rhythm (Leranth & Kiss, 1996). Despite the implication of these neurons in behavioral regulation, their role in maternal behavior remains to be elucidated, despite the presence of strong maternal activation.

### LSv^Cb+^ neurons control maternal licking

To determine the function of the LSv^Cb+^ neurons activated by maternal care, we used chemogenetic tools to inhibit these cells in sensitized female mice. These mice take care of the pups similarly to mothers just without nursing (Alsina-Llanes et al., 2015). Specifically, lactating dams and sensitized virgins spend an equivalent amount of time licking their pups (Gubernick & Alberts, 1985). The use of double labeling confirmed that only Cb+ neurons exhibited the DREADD response following AAV injection into the LSv. Given the documented variations in licking behavior exhibited by individual subjects (Pedersen et al., 2011), the effects of chemogenetic inhibition were compared to the animal’s control days. On the day CNO was administered, the mothers spent less time licking their pups. However, this decrease was not apparent on the second control day, by which time the CNO had been excreted from the animals. Thus, the effect is reversible. We could not consistently identify which parts of the pups were licked. However, in line with the literature (Gubernick & Alberts, 1983), the licking was mostly anogenital licking of the pups. Silencing LSv^Cb+^ neurons did not affect other maternal behaviors, suggesting these neurons play a role only in this particular behavior. Furthermore, locomotor activity and anxiety- and depression-like behaviors remained unaffected, which argues against an indirect action on maternal behavior (Y. Wang & Lin, 2025). This contrasts with the effects of inhibiting MPOA GABAergic neurons (Dimén et al., 2021) and galaninerg neurons (Wu et al., 2014), which led to a more general inhibition of maternal behaviors. Thus, the LSv does not appear to regulate maternal motivation globally, but rather a specific component: pup licking. Pup licking is far more than just an affectionate contact, as it is an essential physiological caregiving mechanism with several critical functions.

Anogenital licking induces urination and defecation via spinal reflexes (Gubernick & Alberts, 1983) while general licking supports thermoregulation, arousal, and respiration (Moore & Chadwick-dias, 1986). Furthermore, maternal licking provides patterned tactile stimulation that shapes sensory and limbic circuits including genital reflexes (Lenz & Sengelaub, 2009) and modulates hypothalamic-pituitary development. Pups that receive more maternal licking exhibit lower pain sensation (Sakamoto et al., 2021) and lower stress reactivity (Caldji et al., 2000; Champagne et al., 2001). In females, higher maternal licking increases glucocorticoid, oxytocin, and estrogen receptor expression in the MPOA and promotes high-licking maternal phenotypes in adulthood (Liu et al., 1997; Pedersen et al., 2011). More maternal licking enhances oxytocin receptor and estrogen alpha receptor expression in MPOA of female offspring (Peña et al., 2013) ensuring to be high-licking mothers themselves (Mogi et al., 2023), but early weaning causes altered licking behavior after they become mothers (Mogi et al., 2023; Sakamoto et al., 2021). Licking behavior affects the mother’s physiology as well, including salt-water balance (Gubernick & Alberts, 1983) and the neuroendocrine system promoting growth hormone, prolactin, and oxytocin secretion (Stern, 1986). Licking stimulates immobility, an upright crouching posture, and milk ejection in mothers (Stern & Johnson, 1990) and releases oxytocin in the mother strengthening mother–infant bonding (Nagasawa et al., 2012). All of these show how relevant pup licking is in maternal care, and our results defined LSv^Cb+^ neurons as participants in the circuitry of controlling this behavior.

### Characteristics of LSv^Cb+^ neurons

Biocytin filling revealed previously undescribed morphological features of LSv^Cb⁺^ neurons, which have relatively small, elliptical somata and poorly developed dendritic arbors, similar to calbindin-expressing interneurons in the hippocampus and neocortex (Sik et al., 1995). In case of the thalamus, Cb^+^ neurons are often projecting neurons with thin and long axons and less dendritic arborization (Celio, 1990) resembling LSv^Cb+^ neurons, which were also shown here to be predominantly projecting based on anterograde and retrograde tracing. Indeed, the soma size of MPOA-projecting LSv^Cb+^ neurons was comparable to that measured after biocytin labeling. This contrasts with Cb⁺ projection neurons in regions such as the amygdala, which have larger somata (Legaz et al., 2005) and more elaborate dendritic arborization (McDonald et al., 2018).

The electrophysiological properties of Cb⁺ and Cb⁻ neurons in the LSv had not been previously characterized. Both populations displayed regular-spiking behavior and similar spike waveforms in response to depolarizing current steps. However, LSv^Cb+^ neurons showed lower spike amplitude and rise slope but higher membrane resistance than LSv^Cb-^ neurons. Higher input resistance is typically associated with smaller cell size. These features suggest that Cb⁺ neurons are intrinsically more excitable and can respond robustly to relatively small synaptic inputs, consistent with selective activation by behaviorally salient stimuli such as pups.

Maternal behaviors are induced by hormonal changes during late pregnancy and maintained by pup-derived stimuli (Bridges, 2020), In rodents, pups actively solicit care through tactile, auditory, and hormonal cues (Lapp et al., 2024). The posterior intralaminar nucleus (PIL) in the lateral thalamus (Yang et al., 2025) has been proposed as a major relay of pup-related tactile and auditory information (Dobolyi et al., 2018; Valtcheva et al., 2023). PTH2-expressing PIL neurons show pup-induced c-Fos activation in rats (Cservenák et al., 2013), and here we demonstrate a similar activation in mice, suggesting a conserved role. In rats, PIL PTH2⁺ neurons project to the LSv and target c-Fos–positive LSv neurons (Puska et al., 2025). We further show that pup-activated LSv^Cb⁺^ neurons preferentially receive PTH2 input compared with non-activated Cb⁺ and Cb⁻ cells, supporting the idea that the excitatory (Keller et al., 2022) PIL–LSv PTH2 pathway conveys pup-related signals to LSvCb⁺ neurons. PTH2-containing terminals were also found to innervate GABAergic LSv neurons, and most were located near c-Fos–positive LSv^Cb⁺^ cells. The PTH2 receptor (PTH2R) is abundant in the LS (Dobolyi et al., 2006; Faber et al., 2007), and we show that many LSv^Cb⁺^ neurons in females express PTH2R, further supporting direct excitation of maternally relevant LSv^Cb⁺^ neurons by PIL input.

The mechanisms by which licking influences maternal physiology remain unclear, but the LSv, which regulates licking behavior, is a strong candidate. To fulfill this role, it must project widely to hypothalamic and other maternal control regions. The projection pattern of LSv^Cb⁺^ neurons largely matches that described for the LSv using non–cell-type-specific tracers (Deng et al., 2019; Risold & Swanson, 1997b; Rizzi-Wise & Wang, 2021), although projections to the medial amygdala, habenula, and midline thalamus were weaker, and projections to the PVN and VMH were absent, suggesting these targets are preferentially innervated by LSv^Cb⁻^ neurons. Retrograde tracing from one major target, the MPOA, labeled 91% of LSv neurons, about two-thirds of which were Cb⁺, confirming a prominent LSv^Cb⁺^→MPOA pathway. Given the established role of the MPOA in regulating pup licking (Champagne et al., 2001; Peña et al., 2013), these findings support the view that LSv^Cb⁺^ neurons influence maternal licking via their projections to core maternal control circuits.

*In conclusion*, the present paper provides evidence that LSv^Cb+^ neurons play a role in controlling pup licking behavior. The function of these neurons is likely influenced by inputs from the pups via PTH2 neurons in the PIL. In turn, LSv^Cb+^ neurons project to various brain regions, primarily different hypothalamic regions.

## Conflict of interest statement

The authors declare no conflict of interest.

## Author contribution

Vivien Szendi participated in the design of the experiments, performing surgical procedures and histological work, analyzing behavioral test results, interpretation of the results and literature, and writing of the manuscript. Gina Puska took part in design of the experiments, light and electron microscopic histological work and analyses, interpreting data and writing of the manuscript. Máté Egyed, Miklós Márton Takács and Szilvia Bartók participated in brain sectioning and immunolabeling extensively. Attila Szűcs, Petra Varró and Júlia Puskás performed the electrophysiological recordings including slice preparation, data analysis and interpretation of results. Árpád Dobolyi participated in the design of the experiments, data analyses, interpretation of the results and the literature, acquisition of funds and writing of the manuscript.

## Acknowledgements

This study was supported by the DKOP-23 Doctoral Excellence Program of the Ministry for Culture and Innovation from the source of the National Research, Development and Innovation Fund and Gedeon Richter Excellence PhD Grant funded by the Gedeon Richter Talentum Foundation of Gedeon Richter Plc. (H-1103 Budapest Gyömrői street 19-21.) for VSz. This study was supported by the strategic research fund of the University of Veterinary Medicine Budapest (Grant No. SRF-003) for GP. Support by EKÖP-25-2-41 University Research Scholarship Program of the Ministry for Culture and Innovation from the source of the National Research, Development and Innovation Fund was provided for EM. ASz was supported by the National Research, Development and Innovation Office under grant ANN_135291. Grant support was provided by the National Brain Research Program NAP2022-I-3/2022 (NAP 3.0) and NAP3-KOLL-2024-1/2024 of the Hungarian Academy of Sciences for AD, the National Research, Development and Innovation Office NKFIH OTKA K146077, NKKP Excellence 151425, ERA-NET NEURON JTC 2024 grant of the Ministry of Innovation and Technology of Hungary, as well as CELSA/24/0205 of Eotvos University.

We also appreciate the technical assistance of Nikolett Hanák and Ágnes Konrád.

## Notes

### Competing Interest Statement

The authors have declared no competing interest.

